# Baobab Diversity, Seed Germination and Early Growth

**DOI:** 10.1101/763177

**Authors:** Kenneth F. Egbadzor

## Abstract

The diversity in baobab was studied on 75 trees located at Adaklu District and Ho and Hohoe Municipalities. Thirteen morphological traits were used in the characterisation based on Bioversity descriptors for baobab. GenStat edition 12 was used to analyse the diversity as well as germination and growth data. Group average hierarchical clustering with Jaccard similarity coefficient discriminated among most of the baobab trees. Clustering was not based on location although few trees that were not discriminated were from the same communities. The clustering can be used in selecting trees for further studies and domestication. Germination tests were conducted with soaked, boiled and sulphuric acid treated seeds. Only the sulphuric acid treated seeds had germination significantly higher than the control. More studies should be done to find easier way of breaking seed dormancy of baobab. There were significant differences in 100 seed weight, seed length and thickness among seeds from three different trees. However, there were no significant difference in seed width of the same sample. Seed size traits should be considered in selecting baobab for domestication because of the high variability revealed. Observation on seedling growth revealed less than 10 leaves in the first month and increased to about 20 in the fourth month. Growth in girth was 5mm and 8mm in the first and fourth months respectively. Seedling height of 17cm in the first month reached 40cm in four months. Information from this research is valuable for further work on domestication of baobab.

## Introduction

Baobab (*Adansonia digitata* L.) is an ancient African tree with important uses across the continent (Rashford 2015). It is also gaining international prominence especially in the food, cosmetic and pharmaceutical industries worldwide (Gruenwald and Galizia 2005). Despite its importance, the tree remains in the wild undomesticated. Fortunately, because of its status, there is a growing interest to domesticate it (Jensen *et al.*, 2011). It is considered one of the most important underutilised crops for domestication (Gebauer *et al.*, 2016).

One of the contributing factors to the underutilisation of baobab is its undomesticated state. Its availability is not controlled by man. The tree grows naturally throughout sub-Saharan Africa with higher presence in the savannahs and drier environments (Chadare *et al.*, 2008). As it originates from Africa, diversity abounds in baobab within the continent. These variabilities in African baobab hold prospect for its domestication and improvement (Korbo *et al.*, 2012). Baobab diversity although reported to some extent, it has not been exploited enough for the development of the crop. The Volta Region of Ghana is a typical location where baobab trees are abundant and can be exploited for domestication.

Apart from the diversity that has to be exploited for domestication, there are issues of seed germination and long juvenile state of baobab that have to be considered. The seed coat of baobab is hard causing mechanical dormancy. The dormancy has to be broken before the seed can germinate. Naturally, it takes a long period of time for baobab seed to germinate on itself and when planted without pre-treatment, germination may be lower than 20% (Danthu *et al.*, 1995). Animals feed on baobab pulp in the wild. As they ingest the seed, the pulp is digested however the seed drop in their faeces. The process the seed goes through in the digestive track of the animals help in breaking dormancy. Other smaller organisms including insects such as cotton stainers (*Dysdercus superstitiosus*) feed on the fruit pulp aiding germination of the seed (SCUC 2006). This natural means of germination is not reliable for crop production. It has been found that the use of sulphuric acid results in breaking of dormancy and hence early and uniform germination of baobab seeds (Niang *et al.*, 2015).

There were three main objectives of the study. First, the study aimed as assessing the diversity of baobab in the study area and identify useful trees that could be use in domesticating the plant. In addition, the study examined techniques that could be used in early germinating of baobab seeds and study the juvenal growth of the plant.

## Materials and Methods

Seventy-five baobab trees were randomly selected from Adaklu District and Ho and Hohoe Municipalities and characterised with 14 morphological traits from bioversity descriptors for baobab (Kehlenbeck *et al.*, 2015). The 14 traits are listed in Table 1. At the District or each Municipality, five communities were selected, and from each community five plants were identified for the studies. This resulted in 25 trees from a District/Municipality.

**Table 1:**
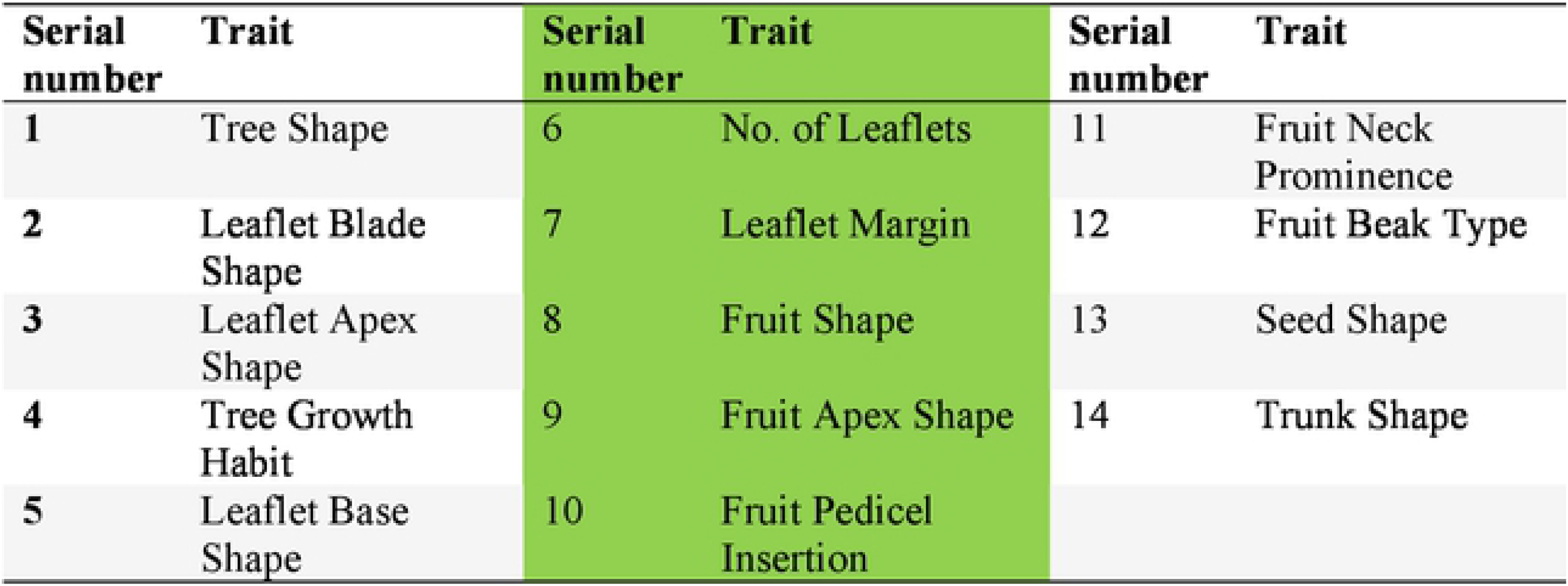
Traits used in characterising baobab accessions

Seeds sizes of three baobab trees were studied. The attributes studied were 100 seed weight, seed length, width and thickness. Electronic scale was used in weighing the 100 seeds. Electronic calliper was used for the seed length, width and thickness measurements as shown in Figures 1, 2 and 3 respectively.

**Figure 1:**
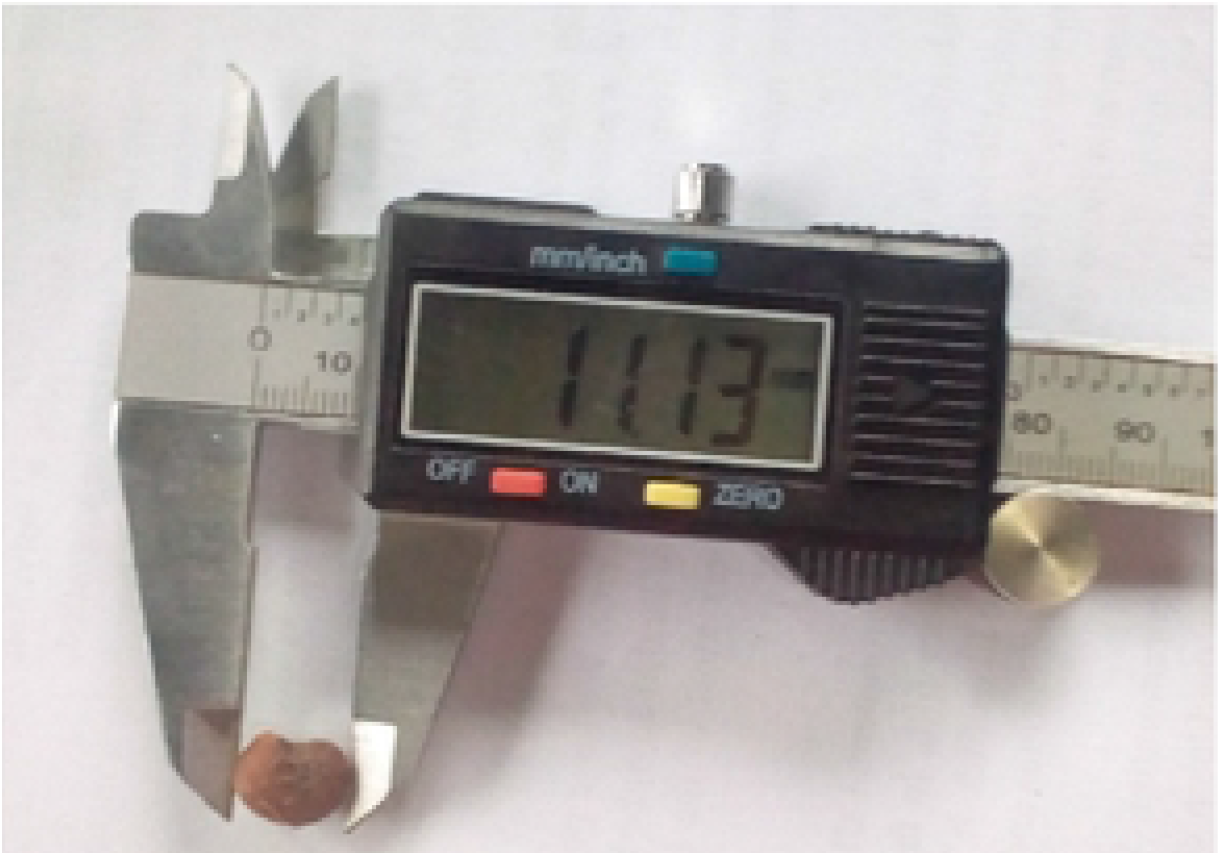
Seed length measurement

**Figure 2:**
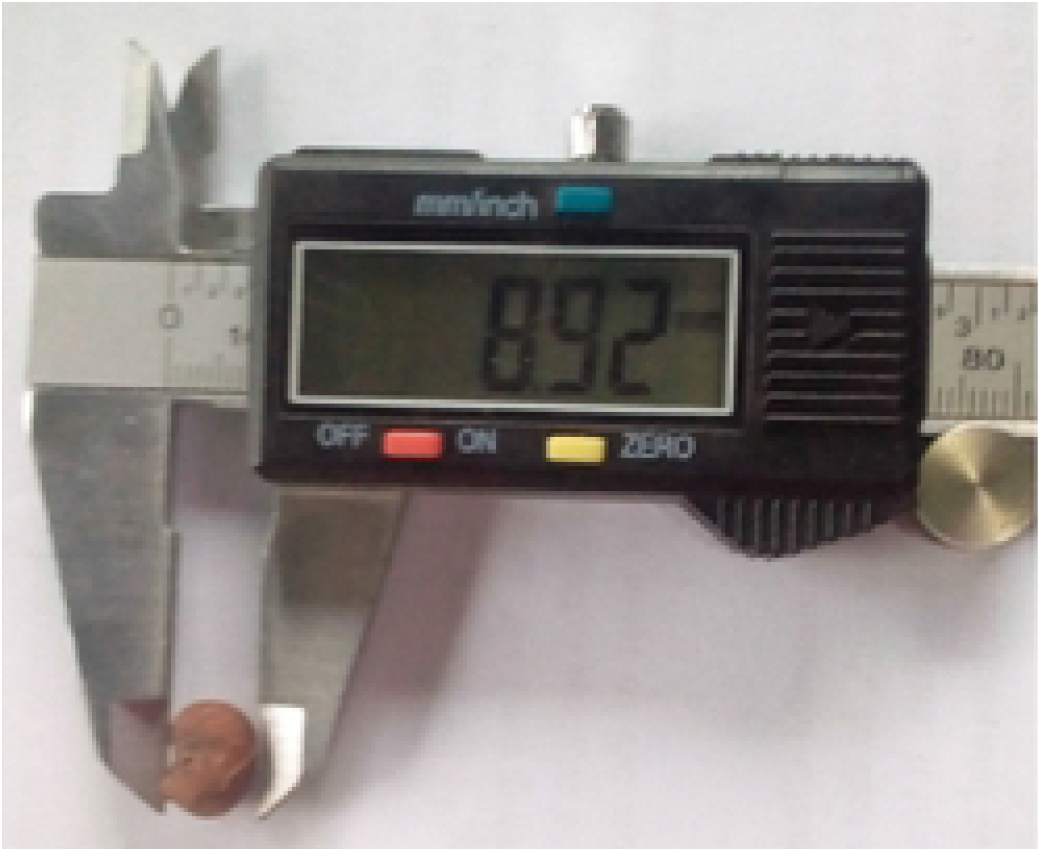
Seed width measurement

**Figure 3:**
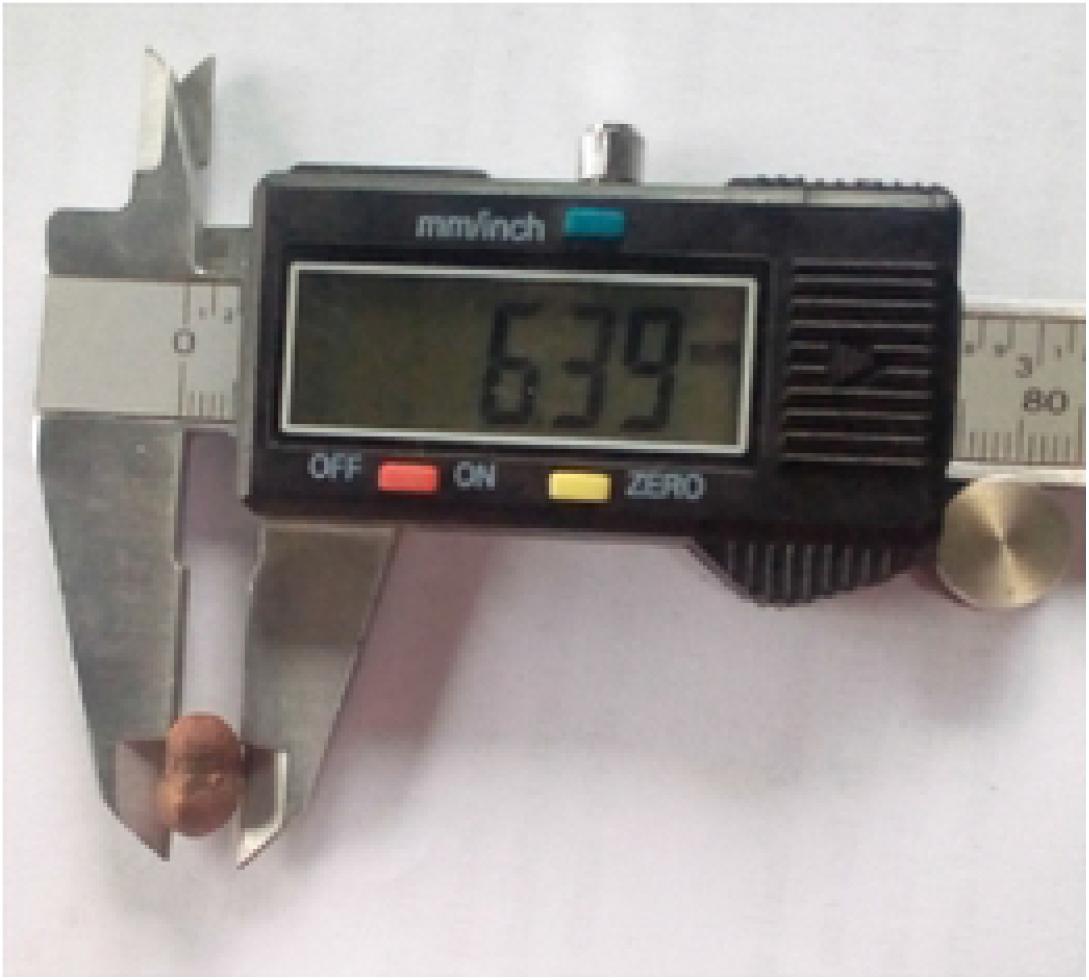
Seed thickness measurement

Germination test was performed using boiling water and sulphuric acid treatments. Seed samples were immersed in boiling water for either two or five seconds. Sulphuric acid treatment involved the immersing of seeds in concentrated sulphuric acid for nine hours. After nine hours, seeds were removed from the acid and washed thoroughly under running tap. Treated seeds were placed on wet paper towels and kept in a dark box.

Another treatment observed was boiling water treatment followed by soaking. After the five seconds immersion in the boiling water, seed samples were soaked in water for twelve to forty-eight hours. After soaking, seeds were placed on wet paper towels and kept in dark boxes for germination.

Growth of baobab seedlings were also observed for four months. Seedlings were examined for growth in number of leaves, girth at soil level and plant height.

### Data analyses

All the data were analysed with GenStat (Payne *et al.*, 2009). Morphological data scored were converted into present or absent data and the dissimilarity used to generate a dendrogram using arithmetic mean method. Analysis of variance was conducted for the data scored on seed size and germination tests. Growth rate was represented on a graph.

## Results

### Diversity studies

Seventy-five trees from Adaklu District and Ho and Hohoe Municipalities were characterised using morphological traits. Phylogenic relationship of the collection is shown in a dendogram in Figure 4.

**Figure 4:**
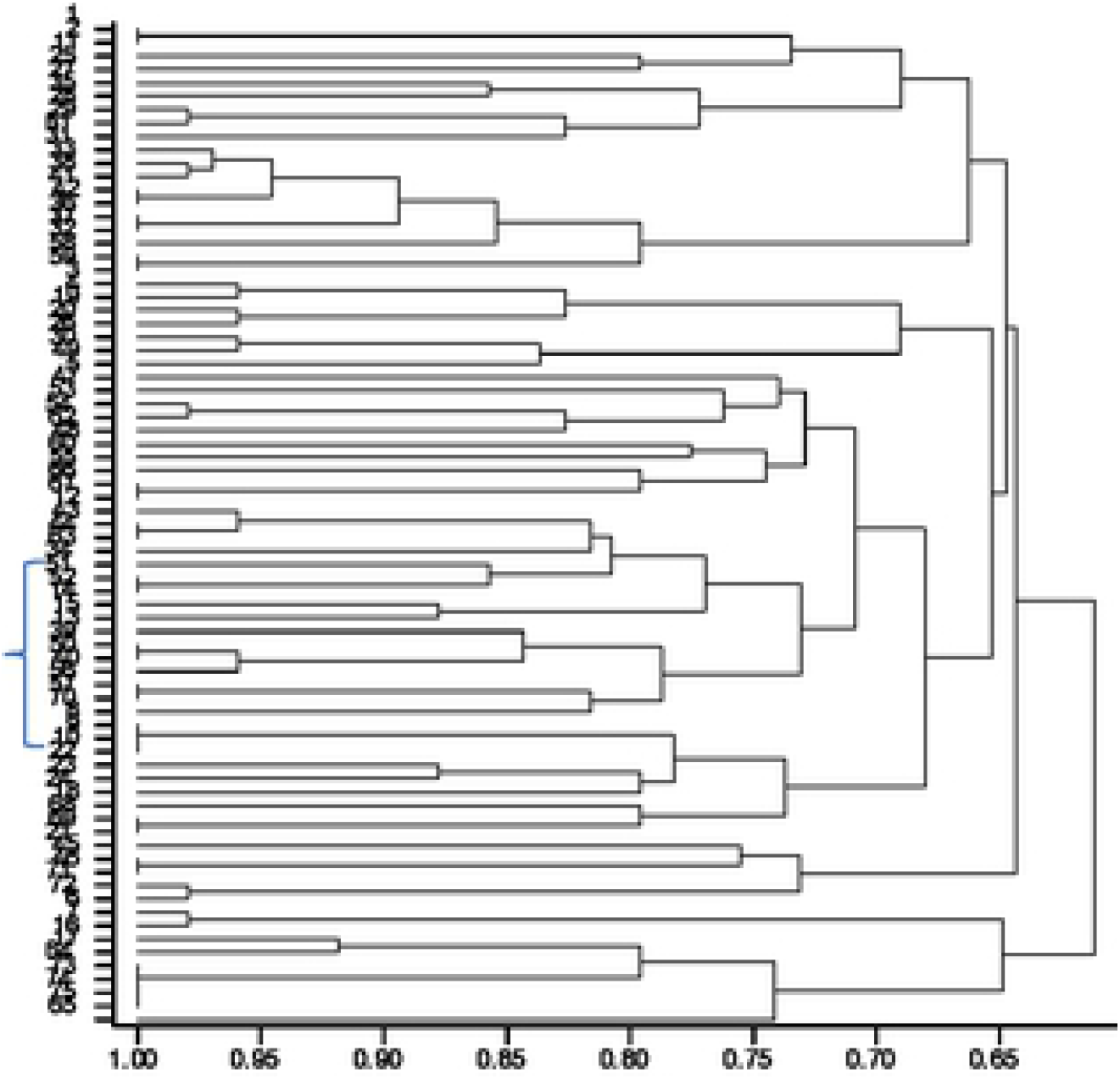
Group average hierarchical clustering of 75 baobab trees based on 13 morphological traits used to calculate Jaccard similarity coefficient

### Seeds Sizes

Seeds from three different trees were analysed for weight, length, width and thickness. The arithmetic means of the seed sizes are summarized in Table 2.

**Table 2.** Sizes of baobab seeds from different trees

| Tree | 100 Seed Weight (g) | Length (mm) | Width (mm) | Thickness (mm) |
| --- | --- | --- | --- | --- |
| One | 42.70 | 11.72 | 8.38 | 6.467 |
| Two | 41.20 | 10.92 | 8.39 | 6.625 |
| Three | 43.50 | 11.18 | 8.67 | 7.159 |
| F-probability | < 0.001 | 0.029 | 0.454 | 0.010 |
| Lsd | 0.575 | 0.583 |  | 0.4417 |

### Germination tests

Sampled seeds from three different trees with different treatments were used for germination test. Seeds were immersed in boiling water for two or five seconds and another immersed in concentrated sulphuric acid for nine hours. Records on germination were taken from the fourth to the twenty-third day after treatment. The results are presented in Table 3.

**Table 3:** Germination of baobab seeds from different trees with different treatments *Note: n = 10 with 4 replications for each treatment*

| Tree | Treatment | Mean Number / % germinations |  |  |  |
| --- | --- | --- | --- | --- | --- |
|  |  | Day 4 | Day10 | Day 14 | Day 23 |
| 1 | Control | 0.0 | 0.25 (2.5) | 0.25 (2.5) | 0.25 (2.5) |
| 1 | 2Sec | 0.0 | 0.0 | 0.0 | 0.0 |
| 1 | 5Sec | 0.0 | 0.0 | 0.0 | 0.0 |
| 1 | H<sub>2</sub>SO<sub>4</sub> | 2.0 (20) | 6.0 (60) | 6.25 (62.5) | 7.00 (70) |
| 2 | Control | 0.25 (2.5) | 0.5 (5) | 0.5 (5) | 0.5 (5) |
| 2 | 2Sec | 0.0 | 0.0 | 0.0 | 0.25 (2.5) |
| 2 | 5Sec | 0.0 | 0.25 (2.5) | 0.25 (2.5) | 1.00 (10) |
| 2 | H<sub>2</sub>SO<sub>4</sub> | 0.5 (5) | 5.25 (52.5) | 6.00 (60) | 6.50 (65) |
| 3 | Control | 0.0 | 0.0 | 0.50 (5) | 1.25 (12.5) |
| 3 | 2Sec | 0.0 | 0.0 | 0.25 (2.5) | 0.75 (7.5) |
| 3 | 5Sec | 0.0 | 0.0 | 0.0 | 0.25 (2.5) |
| 3 | H<sub>2</sub>SO<sub>4</sub> | 1.5 (15) | 3.75 (37.5) | 4.75 (47.5) | 5.75 (57.5) |
| F probability |  | <.001 | <.001 | <.001 | <.001 |
| Lsd |  | 0.8123 | 1.33 | 1.206 | 1.504 |

### Boiling water treatment

Seeds from three different trees were dipped in boiling water for five seconds. Results are presented in Table 4.

**Table 4:** Mean number of germination after boiling water treatment

| Tree | Number of germinations | Percentage germination |
| --- | --- | --- |
| One | 0.50 | 1.25 |
| Two | 1.00 | 2.5 |
| Three | 1.00 | 2.5 |
| F-Probability | 0.128 |  |

### Germination test for boiled and soaked seeds

Seeds dipped in boiling water for five seconds were soaked in water for different time durations and tested for germination. The results are presented in Table 5.

**Table 5:** Germination of boiled and soaked seeds

| Treatment | Number of germinations | Percentage germination |
| --- | --- | --- |
| Soaked for 12hr | 0.25 | 0.6 |
| Soaked for 24hr | 0.25 | 0.6 |
| Soaked for 36hr | 0.50 | 1.25 |
| Soaked for 48hr | 1.00 | 2.5 |
| F-Probability | 0.152 |  |

### Seedling growth rate

Growth of seedlings of baobab in terms of number of leaves, girth and height are represented in Figure 5. Positive correlations were observed between the traits measured. Figures 6 and 7 show baobab seedlings.

**Figure 5:**
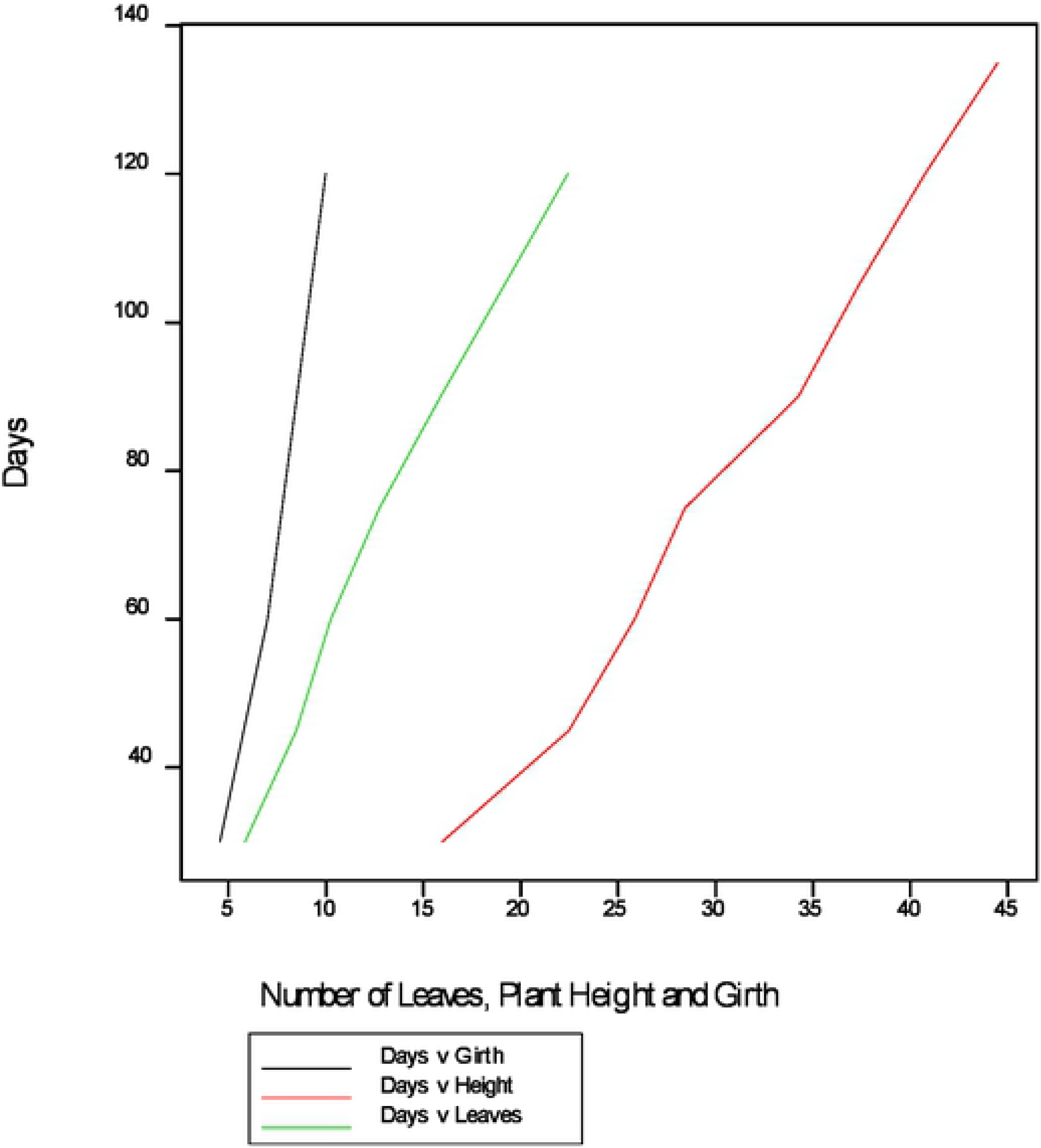
Growth in Number of Leaves, Plant Height and Girth

**Figure 6:**
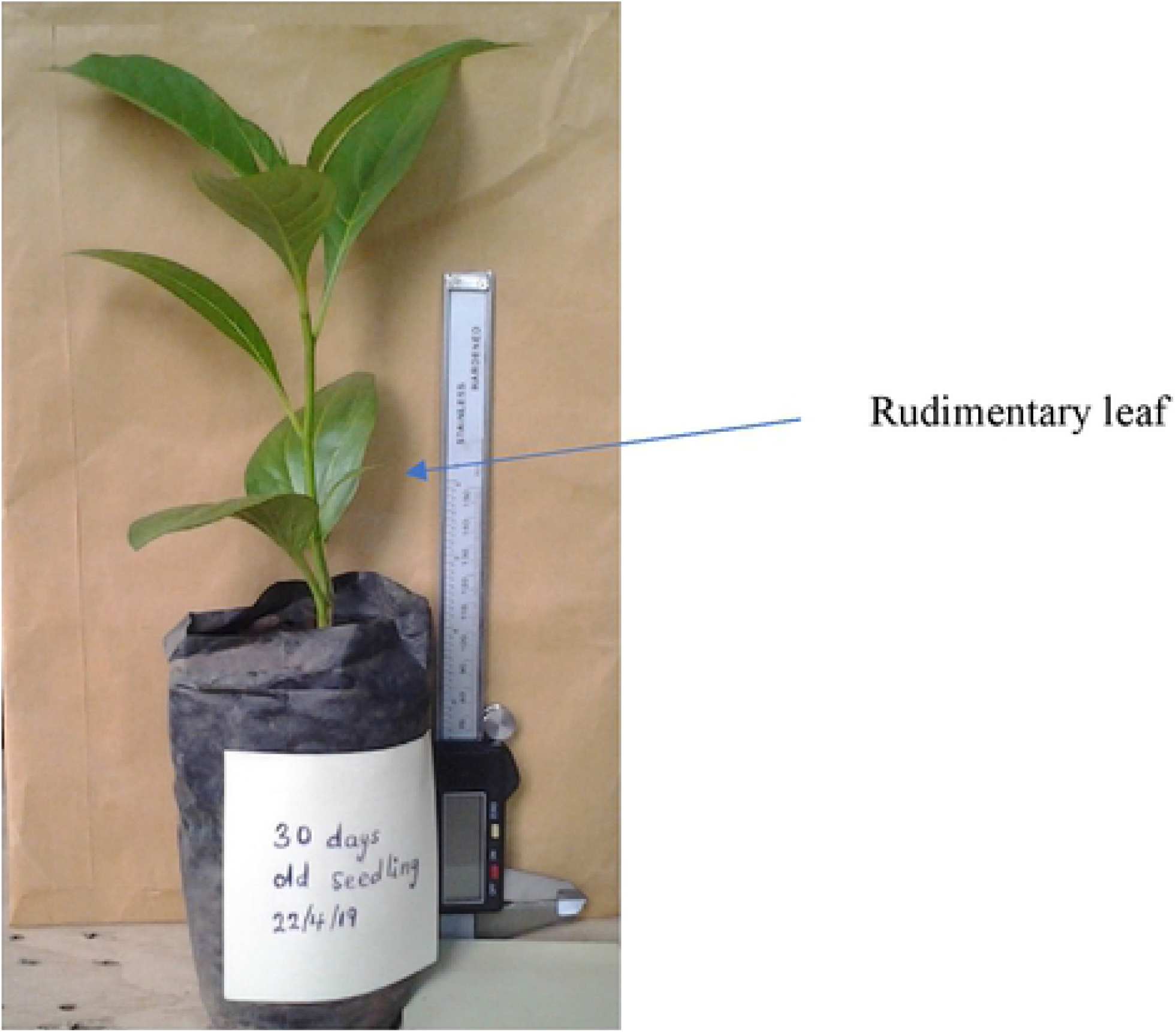
A month old baobab seedling

**Figure 7:**
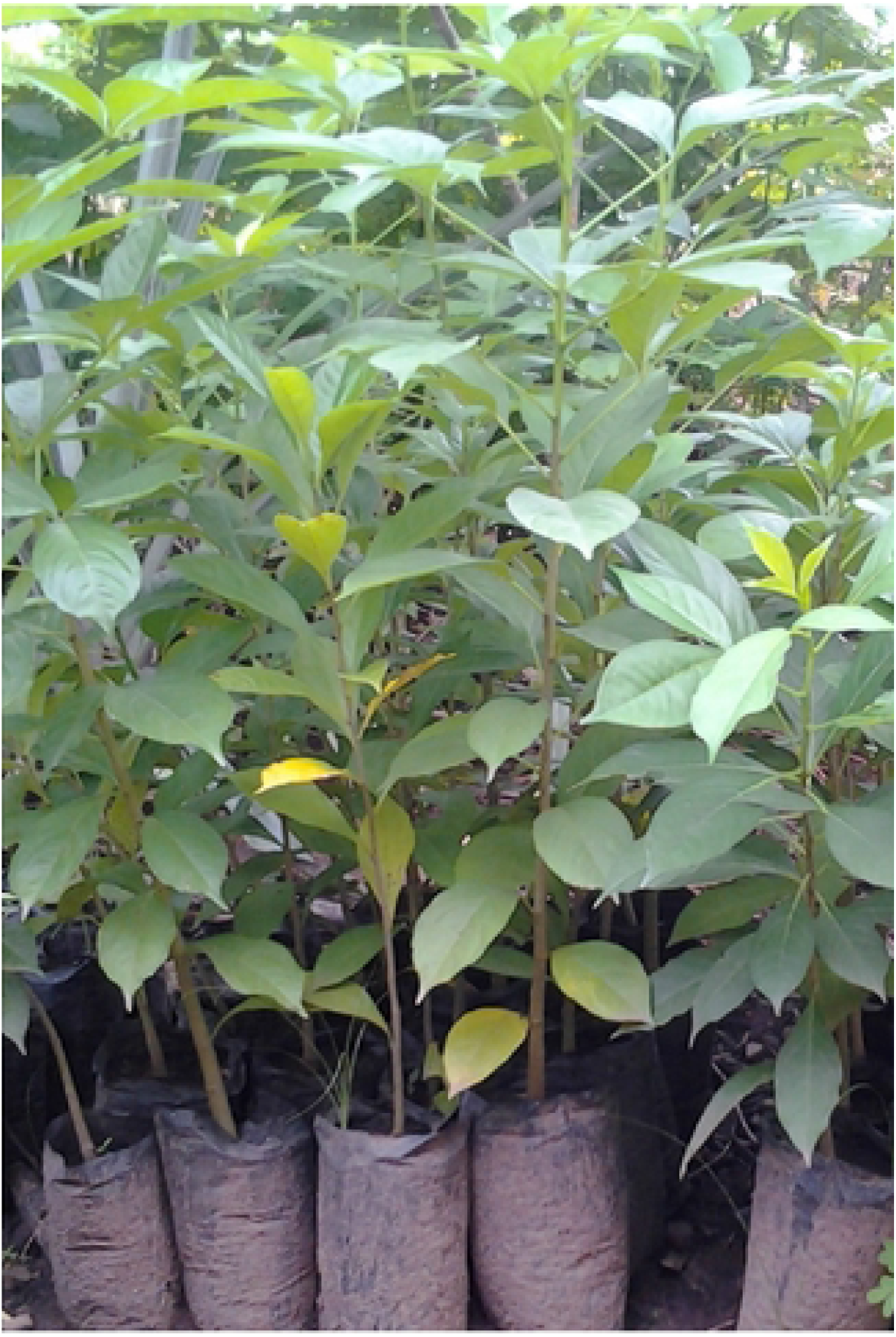
Three Months Old Baobab Seedlings

## Discussion

There were variabilities in all the thirteen morphological traits used to characterise the baobab accessions. For example, fruit shapes observed were obovate, ellipsoid, ovate and oblong from different trees. The traits discriminated among most of the 75 baobab trees. This shows high diversity that exist among the baobab population studied. Assogbadjo *et al* (2010) also found within and between population variability in their study on baobab diversity in Benin, although, the traits they studied were mostly quantitative compared with the qualitative traits in this study. However, despite the variability observed, some of the accessions in the current study could not be separated based on the traits scored. Addition of more traits especially quantitative traits could discriminate among more of the trees. Fruit traits including pulp nutrient content and taste should be considered in further studies because of their importance to consumers of baobab as suggested by Simbo *et al.* (2012).

Clustering generally was not based on locations as trees from different communities, district or municipalities clustered together. Two major groups were formed at 60% similarity coefficient. The smaller group at this level had only nine members with four and seven respectively from Ho and Hohoe Municipalities. A group at 70% similarity had members from the two municipalities and the Adaklu District. Although, clear clustering could not be observed based on location, those trees which could not be separated were mainly from the same localities. These pairs; 1 and 2, 36 and 37 and 73 and 74 collected from Hohoe, Adaklu and Ho respectively did not separate from each other. These were trees sampled from the same communities in the same district or municipalities. It is likely that they are closely related by ancestry. Sampling should be made from different environments to capture wider variability.

Variability was observed in 100 seed weight, seed length and seed thickness for seeds from three different trees. Seed width on the other hand were not significantly different from each other. Inclusion of seed and other quantitative traits could have revealed more diversity within the collection. Seed size of baobab can be explained by seed length and thickness based on the results of this study. These two traits apart from being significantly different for the three samples, they positively correlate as well. Bigger seed size and more pulp also result in higher weight. These traits can be considered in domestication as suggested by Jensen *et al.* (2011). Bigger seeds may be preferred especially when the seed is consumed as a whole nut and may also result in higher oil content. Oil content of the baobab seed is increasingly becoming important in many sectors including, cosmetics, pharmaceutical and food (Komane *et al.*, 2017). Seed traits are therefore essential to study with the aim of domesticating baobab.

Seed germination is important in cultivated crops. Baobab has hard seed coat causing mechanical dormancy that must be broken before germination. Naturally, seed dormancy of baobab is broken as they pass through the digestive track of animals (Gruenwald and Galizia 2005). Smaller organisms like cotton strainers may also have effect on germination of baobab as they feed on the pulp.

All the treatments had significantly low germination compared with a nine-hour sulphuric acid treatment. Niang *et al.* (2015) had similar observation with sulphuric acid treatment. Germination started four days after treatment for observation that lasted for twenty-three days. Germination was influenced by both sulphuric acid treatment and the tree from which seeds were collected from. Hot water treatment and control had significantly lower germination than sulphuric acid treatment across the three seed samples. However, tree one had significantly more germinations that tree two and three seeds with sulphuric acid treatment. This suggested that seeds were more viable for tree number one than the other two trees. The germination could have been better than the 70% record if the seeds were more viable. Most of the germinations also took place within ten days of the treatment.

Dipping of seeds in boiling water for up to five seconds did not result in germination. There was more intense treatment by soaking boiled seeds for up to forty-eight hours. This also did not result in significant germination of baobab seeds. We tried boiling the baobab seeds between two and ten minutes based on the report of SCUC (2006). However, our results were only rots. Danthu *et al* (1995) reported that response to boiling water varied from a seed lot to the other. Therefore, the boiling water treatment can be tried for seeds from different trees. Growth of baobab seedlings were observed for four months. One and three month old seedlings are shown in Figure 6 and 7 respectively. Baobab is a dicotyledonous plant with hypogeal germination. It has a pair of leaves initially and then a rudimentary one shown in Figure 6. It then develops a series of alternating simple leaves with short petioles. After about ten simple leaves, it develops compound leaves with lobes on long petioles. It commonly starts with three lobes and increases to five. The leaf lobe can be entire or undulating.

Growth in number of leaves, height and girth is slow for baobab. For the first month, baobab developed eight or less number of leaves. Thereafter, one leaf on the average is developed every two weeks.

Growth in girth was about 8mm in three months. This shows slow initial growth of baobab. Growth in height was less than 20cm in the first month. After four months, growth in height reached over 40cm.

## Conclusion

There is high diversity in baobab based on morphology in the Volta Region of Ghana. The diversity can be useful in selecting plants for domestication. It is suggested that end user traits like taste of baobab fruit pulp should be included in characterisation. From the studies, sulphuric acid treatment is the only reliable means to quickly and uniformly germinate baobab seeds. Boiling water treatment reported by some authors did not work in our case. Less expensive and reliable methods need to be investigated. Early growth of baobab is slow in terms of plant height, number of leaves and growth in girth. The study has laid foundation for further studies and domestication of baobab.

## Acknowledgment

I appreciate all those who are encouraging me to study baobab. Special thanks to my student Mr. Pius Gator who helped in data collection.

